# MinION™, a portable long-read sequencer, enables rapid vaginal microbiota analysis in a clinical setting

**DOI:** 10.1101/2021.01.07.425663

**Authors:** Shinnosuke Komiya, Yoshiyuki Matsuo, So Nakagawa, Yoshiharu Morimoto, Kirill Kryukov, Hidetaka Okada, Kiichi Hirota

**Affiliations:** Department of Obstetrics and Gynecology, Kansai Medical University Graduate School of Medicine, Osaka, Japan; HORAC Grand Front Osaka Clinic, Osaka, Japan; Department of Human Stress Response Science, Institute of Biomedical Science, Kansai Medical University, Osaka, Japan; Department of Molecular Life Science, Tokai University School of Medicine, Kanagawa, Japan; Department of Genomics and Evolutionary Biology, National Institute of Genetics, Shizuoka, Japan

**Keywords:** 16S rRNA, bacterial vaginosis, long-read sequencer, MinION™, nanopore

## Abstract

It has been suggested that the local microbiota in the reproductive organs is relevant to women’s health and may also affect pregnancy outcomes. Analysis of partial 16S ribosomal RNA (rRNA) gene sequences generated by short-read sequencers has been used to identify vaginal and endometrial microbiota, but it requires a long time to obtain the results, making it unsuitable for the rapid analysis of small samples in a clinical context. We demonstrated a simple workflow using the nanopore sequencer MinION™ that allows high-resolution and rapid differentiation of vaginal microbiota. Vaginal samples collected from 18 participants were subjected to DNA extraction and full-length 16S rRNA gene sequencing with MinION™. The principal coordinate analysis showed no differences in the bacterial compositions regardless of the sample collection method. The vaginal microbiota results could be reported within 2 days of specimen receipt. Although bacterial vaginosis (BV) was not diagnosed by the Nugent score in any cases, groups with both healthy and BV-like vaginal microbiota were clearly characterized by MinION™ sequencing. We conclude that full-length 16S rRNA gene sequencing analysis with MinION™ provides a rapid means for identifying vaginal bacteria with higher resolution. Species-level profiling of human vaginal microbiota by MinION™ sequencing can allow the analysis of associations with conditions such as genital infections, endometritis, and threatened miscarriage.

## Introduction

Bacterial vaginosis (BV) is one of the most common gynecological disorders among women of reproductive age [1]. It is caused by disruption of the vaginal microbiota due to various factors such as psychological stress, poor physical condition, menstrual cycle, and pregnancy. It has relatively mild subjective symptoms, such as abnormal vaginal discharge (odor, color, amount of discharge) and itching, but there are cases of unrecognized infection and recurrent infections without significant symptoms [1]. BV has been clinically reported to cause adverse reproductive outcomes, including sexually transmitted infections, genitourinary viral infections such as HIV-1/2, HPV, and HSV-2 [2], and preterm delivery [3].

Currently, Amsel’s criteria based on clinical outcomes [4] and the Nugent score using Gram stain findings [5] are used as diagnostic criteria for BV. These methods are easy to use, but since they are based on the subjective opinion of the physician, the diagnosis largely depends on the skill of the examiner [6]. Bacterial culture of vaginal discharge is also a standard method to investigate the vaginal microbiota, but some bacteria are difficult to culture [6]. Moreover, in routine gynecological practice, vaginal discharge is not sampled unless patients have symptoms suggestive of vaginosis. Therefore, accurately assessing the vaginal microbiota has been difficult with these regular clinical methods. Due to the lack of a reliable methodology, the association of BV with other vaginal disorders has not been fully elucidated. Until recently, a more detailed analysis of vaginal microbiota in asymptomatic women was considered excessive and might not be clinically relevant. Given that BV has been implicated in women’s health outcomes in various ways, it is crucial to develop a rapid and accurate means to diagnose BV with higher sensitivity. The increased risk of various infections, infertility, miscarriage and preterm birth due to BV has been shown to be impacted by the species that make up the vaginal microbiota [1–3]. In this context, accurate representation of the vaginal microbiota will allow for more appropriate medical treatment. If abnormal vaginal microbiota is identified with persistent cervical lesions, the dysbiosis should be treated even in the absence of symptoms. Detection of asymptomatic BV will potentially be beneficial for the diagnosis and prevention of gynecological diseases.

16S ribosomal RNA (rRNA) gene sequencing analysis using next-generation sequencers can be used to identify bacteria without culture in a manner not dependent on subjective assessment by the examiner. Previous studies have shown that the vaginal microbiota of healthy women is classified into community state type (CST) I to V [7]. CST is defined by the dominant bacteria as follows: CST I: *Lactobacillus crispatus*, CST II: *Lactobacillus gasseri*, CST III: *Lactobacillus iners*, CST IV: lack of *Lactobacillus* spp., and CST V: *Lactobacillus jensenni*. These narrow definitions mean that undiagnosed cases of vaginal dysbiosis may be present. This new classification of the vaginal microbiota has been clarified by performing sequencing analysis on cases with no clinical symptoms. Because a deeper and more accurate understanding of the vaginal microbiota may provide new clinical insights, we should demonstrate that 16S rRNA analysis is quick and easy to perform in a broad range of contexts in clinical practice. MinION™ detects the signal of a DNA nucleotide that passes through a nanopore arranged on a flow cell [8]. This long-read sequencing technology can be used to identify bacterial microbiota with the whole region of the 16S rRNA. MinION™ is smaller, lighter, and easier to deploy than conventional massively parallel sequencing platforms, and it also offers the advantage of high-resolution microbiota analysis with long-read sequencing [9–11]. We constructed an in-hospital vaginal microbiota-analyzing workflow using MinION™. This is the first report to demonstrate that MinION™ can provide rapid patient feedback regarding vaginal microbiota analysis.

## Materials and methods

### Ethics statement

The study adhered to the Declaration of Helsinki throughout the protocol, and written informed consent was obtained from all participants. This study is a cross-sectional observational trial approved by the institutional review board of the medical corporation Sankeikai (Approval Number: 2019-38). After the approval, the study protocol was submitted to the UMIN-CTR clinical trial registry site (https://www.umin.ac.jp/ctr/index.htm) (trial registration number: UMIN000038032), which met the criteria of the International Committee of Medical Journal Editors (ICMJE).

### Recruitment of participants

From April 2019 to May 2020, participants who met all of the following enrollment criteria were recruited: 1) primary infertility patients attending HORAC Grand Front Osaka Clinic in Osaka, Japan; 2) premenopausal Japanese women who were not pregnant; 3) no clinical symptoms strongly suggestive of BV; 4) scheduled hormone replacement embryo transfer cycle with frozen-thawed blastocyst; and 5) those who provided written consent. The exclusion criteria were as follows: 1) known history of HIV-1/2, hepatitis B/C, syphilis, or genital chlamydia infection; 2) known history of diabetes; 3) known history of cervical/vaginal surgery; 4) known history of intrauterine device use; and 5) known history of antibiotic, steroid, or vaginal suppository use within 2 weeks. In addition to collecting vaginal samples, details of age, body mass index, and basal hormone levels (anti-Mullerian hormone, luteinizing hormone, and follicle-stimulating hormone on days 2–4 of the menstrual cycle), medical history, and infertility treatment history were collected from the medical records. All vaginal samples were collected on the day of embryo transfer.

### Nugent score

Vaginal smears were analyzed by two pathologists affiliated with an external laboratory, and the Nugent score was calculated based on microscopic findings: the numbers of *Lactobacillus* (scored as 0 to 4), *Gardnerella* (scored as 0 to 4), and *Mobiluncus* (scored as 0 to 2). A total score of 0 to 3 was diagnosed as representative of healthy vaginal microbiota, 4 to 6 as an intermediate group, and 7 or more as BV [5]. Inappropriate samples with defective specimen collection were diagnosed as indeterminate. If BV was diagnosed based on the Nugent score, our policy was to treat it using antibiotics.

### Vaginal sample collection method

Vaginal lavage samples were collected from 18 participants; from four participants, additional swab samples were collected with written consent. When both lavage and swab samples were collected, the swab samples were collected first. For the lavage method, the inside of the vagina was washed with 10 ml of sterile saline, and at least 2 ml was collected using a sterile syringe. Collected lavage samples were stored at −30 °C until DNA extraction. For the swab method, OMNIgene vaginal kit (OMR-130; DNA Genotek Inc., Ottawa, Canada) was used to infiltrate vaginal mucus from the uterovaginal region, and the swab tip was stored in the preservative solution of the sampling kit. Swab samples were stored at room temperature until DNA extraction in accordance with the manufacturer’s instructions.

### Sequencing sample preparation

DNA was extracted from 22 human vaginal samples using the QIAamp UCP Pathogen Mini Kit (QIAGEN, Venlo, Netherlands) following the manufacturer’s instructions. Briefly, samples were subjected to mechanical cell lysis by bead-beating [12], and DNA was isolated using silica membrane-based spin columns. After extraction, the DNA content was measured using a NanoDrop^®^ 1000 Spectrophotometer (Thermo Fisher Scientific, MA, USA), and the concentrations of extracted DNA from the swab and lavage samples were 6.1–32.3 ng/μl and 53.1–1092.4 ng/μl, respectively.

### MinION™ sequencing

As described in our previous report [12], a slightly modified version of four-primer polymerase chain reaction (PCR) with rapid adapter attachment was performed (https://dx.doi.org/10.17504/protocols.io.bs9knh4w). For amplification of the V1–9 region of the 16S rRNA gene, a forward primer (S-D-Bact-0008-c-S-20) with the anchor sequence 5’-TTTCTGTTGGTGCTGATATTGCAGRGTTYGATYMTGGCTCAG-3’ and a reverse primer with the anchor sequence 5’-ACTTGCCTGTCGCTCTATCTTCCGGYTACCTTGTTACGACTT-3’ were used as inner primers. For amplification of the V3–4 region, 341F with the anchor sequence 5’-TTTCTGTTGGTGCTGATATTGCCCTACGGGNGGCWGCAG-3’ and 806R with the anchor sequence 5’-ACTTGCCTGTCGCTCTATCTTCGGACTACHVGGGTWTCTAAT-3’ were used as inner primers. PCR amplification of the 16S rRNA gene was performed using the KAPA2G™ Robust HotStart ReadyMix PCR Kit (Kapa Biosystems, MA, USA) with an inner primer pair (50 nM for each) and the outer primer mixture (10 nM) of PCR Barcoding Kit (SQK-PBK004; Oxford Nanopore Technologies, Oxford, UK) in a total volume of 25 μl. Each sample was assigned an individual barcode, which was used to identify the PCR product. Amplification was performed under the following PCR conditions: initial denaturation at 95 °C for 3 min; five cycles of 15 s at 95 °C, 15 s at 55 °C, and 30 s at 72 °C; and 30 cycles of 15 s at 95 °C, 15 s at 62 °C, and 30 s at 72 °C; followed by a final extension at 72 °C for 1 min. The amplification products were confirmed using 1% agarose gel electrophoresis (1 × TAE buffer) to be approximately 1600 base pairs (V1–9 region) and 400 base pairs (V3–4 region), which are the expected amplification sizes. After confirmation, the amplified DNA was purified using AMPure^®^ XP (Beckman Coulter, CA, USA). A total of 100 ng (10 μl) of purified DNA was incubated with 1 μl of Rapid Adapter at room temperature for 5 min. The concentrations of swab samples and lavage samples were 8.3–34 ng/μl and 3.58–61 ng/μl, respectively, as determined using a Quantus™ Fluorometer (Promega, WI, USA). The prepared DNA library (total of 11 μl) was mixed with 34 μl of Sequencing Buffer, 25.5 μl of Loading Beads, and 4.5 μl of water. The final adjusted sample was loaded into the flow cell R.9.4.1 (FLO-MIN106; Oxford Nanopore Technologies Ltd.) and attached to the MinION™ Mk1B, which was connected to a personal computer. In this study, four to six samples were loaded at one time and each sequencing session lasted about 90 min. DNA sequencing data were acquired in FAST5 format using MINKNOW software ver. 1.11.5 (Oxford Nanopore Technologies Ltd.).

### Bioinformatic analysis workflow

We performed the bioinformatic analysis using a pipeline of multiple programs built in accordance with our previous reports [9,11]. An overview is given below.

1. Computer: Apple iMac 27-inch, Late 2015 (OS, macOS 10.14.6; CPU, 3.3 GHz Intel Core i5-6600; memory, 16 GB)
2. GUPPY software ver. 3.1.5 (Oxford Nanopore Technologies Ltd.): base calling, conversion of FAST5 files to FASTQ files with quality control based on quality score > 7.
3. SeqKit software ver. 0.10.0 [13]: extraction of a read length of 1,300 to 1,950 bp, based on the 16S rRNA length registered in the SILVA rRNA database ver. 132 (https://www.arb-silva.de/) [14].
4. TANTAN program ver. 18 [15]: removing simple repeat sequences.
5. Minimap2 program ver. 2.14 [16]: eliminating human genome information (Human Genome Assembly GRCh38, https://www.ncbi.nlm.nih.gov/assembly/GCF_000001405.26/) and matching each genome read to the 5,850 representative bacterial genome sequences (Table S1) stored in the GenomeSync database (http://genomesync.org) [11,12].
6. In-house Perl scripts (Genome Search Toolkit, http://kirill-kryukov.com/study/tools/gstk/): selecting the species with the highest Minimap2 score and determining the taxa based on the NCBI taxonomy database.
7. Krona software version 2.7 [17]: visualizing the frequency of detected bacterial species in a given sample.

### Statistical analysis

Principal coordinate analysis was performed and the results were visualized with R version 4.0.2.

## Results

### Characteristics of participants

Our study consisted of 18 Japanese, ranging from 30 to 43 years of age, with a median age of 36.5. All participants had primary infertility with normal menstrual cycles, along with no underlying medical conditions (Table 1).

**Table 1.**
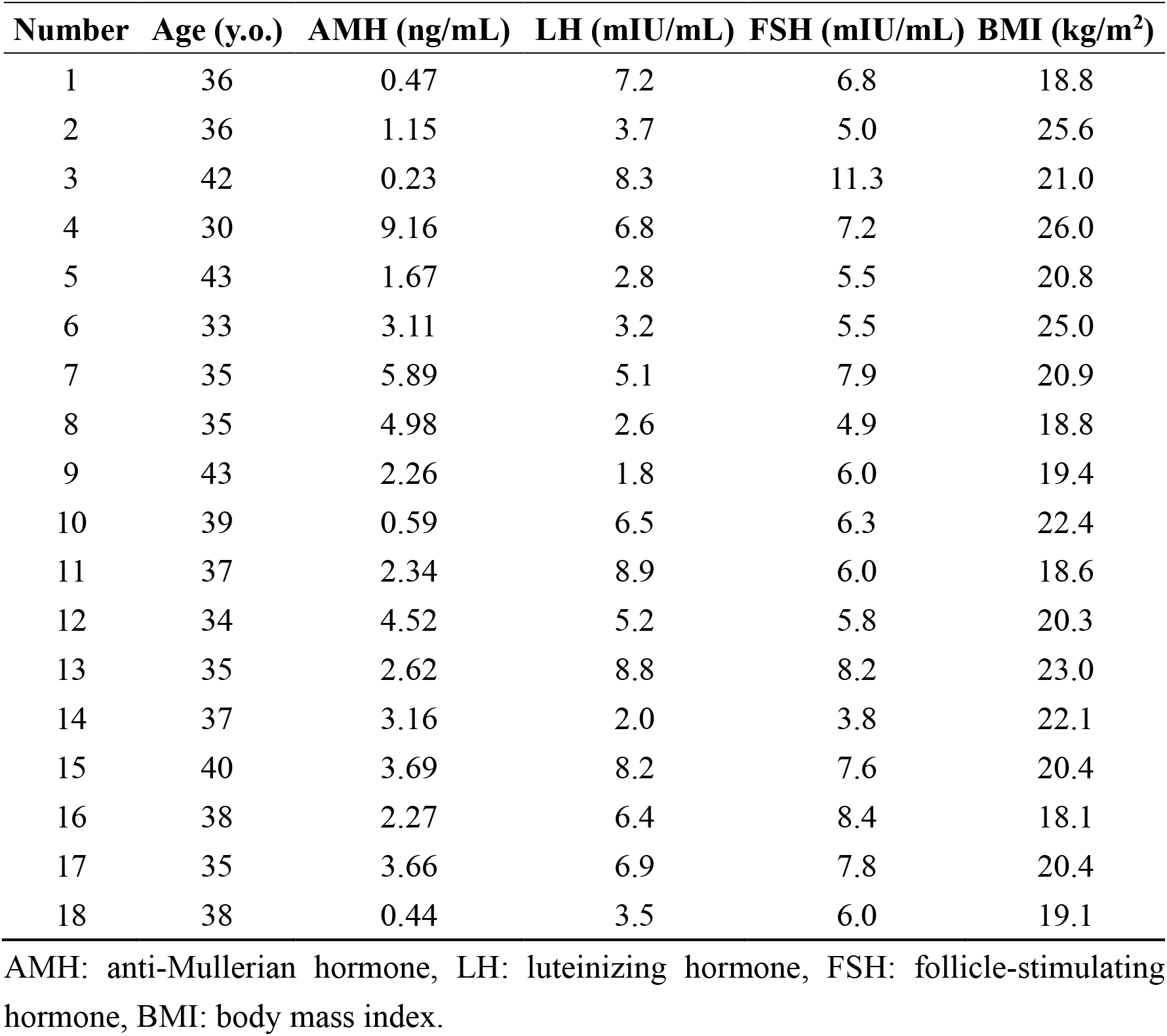
Participants’ background.

| Number | Age (y.o.) | AMH (ng/mL) | LH (mIU/mL) | FSH (mIU/mL) | BMI (kg/m<sup>2</sup>) |
| --- | --- | --- | --- | --- | --- |
| 1 | 36 | 0.47 | 7.2 | 6.8 | 18.8 |
| 2 | 36 | 1.15 | 3.7 | 5.0 | 25.6 |
| 3 | 42 | 0.23 | 8.3 | 11.3 | 21.0 |
| 4 | 30 | 9.16 | 6.8 | 7.2 | 26.0 |
| 5 | 43 | 1.67 | 2.8 | 5.5 | 20.8 |
| 6 | 33 | 3.11 | 3.2 | 5.5 | 25.0 |
| 7 | 35 | 5.89 | 5.1 | 7.9 | 20.9 |
| 8 | 35 | 4.98 | 2.6 | 4.9 | 18.8 |
| 9 | 43 | 2.26 | 1.8 | 6.0 | 19.4 |
| 10 | 39 | 0.59 | 6.5 | 6.3 | 22.4 |
| 11 | 37 | 2.34 | 8.9 | 6.0 | 18.6 |
| 12 | 34 | 4.52 | 5.2 | 5.8 | 20.3 |
| 13 | 35 | 2.62 | 8.8 | 8.2 | 23.0 |
| 14 | 37 | 3.16 | 2.0 | 3.8 | 22.1 |
| 15 | 40 | 3.69 | 8.2 | 7.6 | 20.4 |
| 16 | 38 | 2.27 | 6.4 | 8.4 | 18.1 |
| 17 | 35 | 3.66 | 6.9 | 7.8 | 20.4 |
| 18 | 38 | 0.44 | 3.5 | 6.0 | 19.1 |

AMH: anti-Mullerian hormone, LH: luteinizing hormone, FSH: follicle-stimulating hormone, BMI: body mass index.

### Identification of vaginal bacteria

The composition of the vaginal microbiota was investigated by 16S rRNA gene amplicon sequencing on the MinION™ platform (Figure 1). The V1–9 MinION™ sequencing for almost 90 min yielded 15,836–119,745 reads at first and then filtered 10,688–103,212 reads satisfying the conditions of 1,300–1,950 bp and QC > 7 (details in Table 2). The reads were mapped against the GenomeSync reference database for taxonomic assignment. The results of Nugent scoring and microbiota analysis of the filtered 3,000 reads using MinION™ are shown in Table S2. Based on the Nugent score criteria, four out of 18 cases were in the intermediate group, but none of them were diagnosed as BV. The MinION™ sequencing data were also analyzed by an alternative bioinformatics tool. The FASTQ 16S workflow produced almost similar taxonomic profiles (Table S3). This result confirmed the validity of our method for taxonomic classification and the bacterial compositions were comparable regardless of the program and database used.

**Figure 1.**
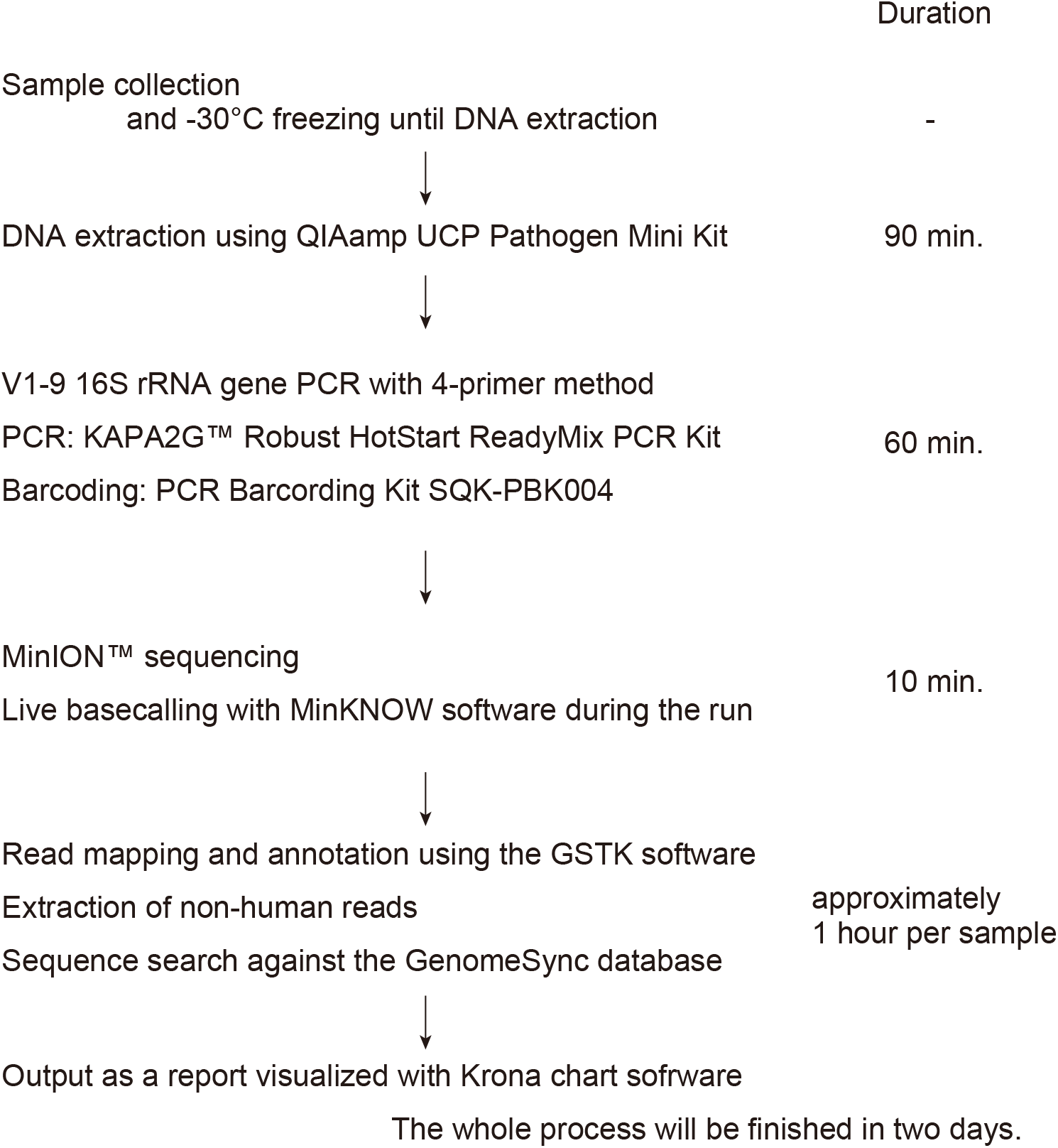
Workflow of 16S rRNA amplicon sequencing with the MinION™ platform and bioinformatic analysis. After collecting the vaginal sample, it should be stored in a freezer at −30°C until DNA extraction begins. Some swab collection kits should be stored at room temperature, in accordance with the manufacturer’s instructions. Sequencing libraries are generated by a four-primer PCR-based strategy; in the initial stages of PCR, the 16S rRNA gene is amplified with an inner primer pair. The PCR product is amplified with the outer primers and targeted to introduce identical barcode and tag sequences at both ends, allowing for the attachment of adapter molecules in a one-step reaction. The library is then loaded into a MinION™ connected to a personal computer. With our experimental method, sequencing runtime of 10 min is sufficient for 3,000 reads to be obtained. A final report of the microbiota can be presented within 2 days of initiating the analysis.

**Table 2.**
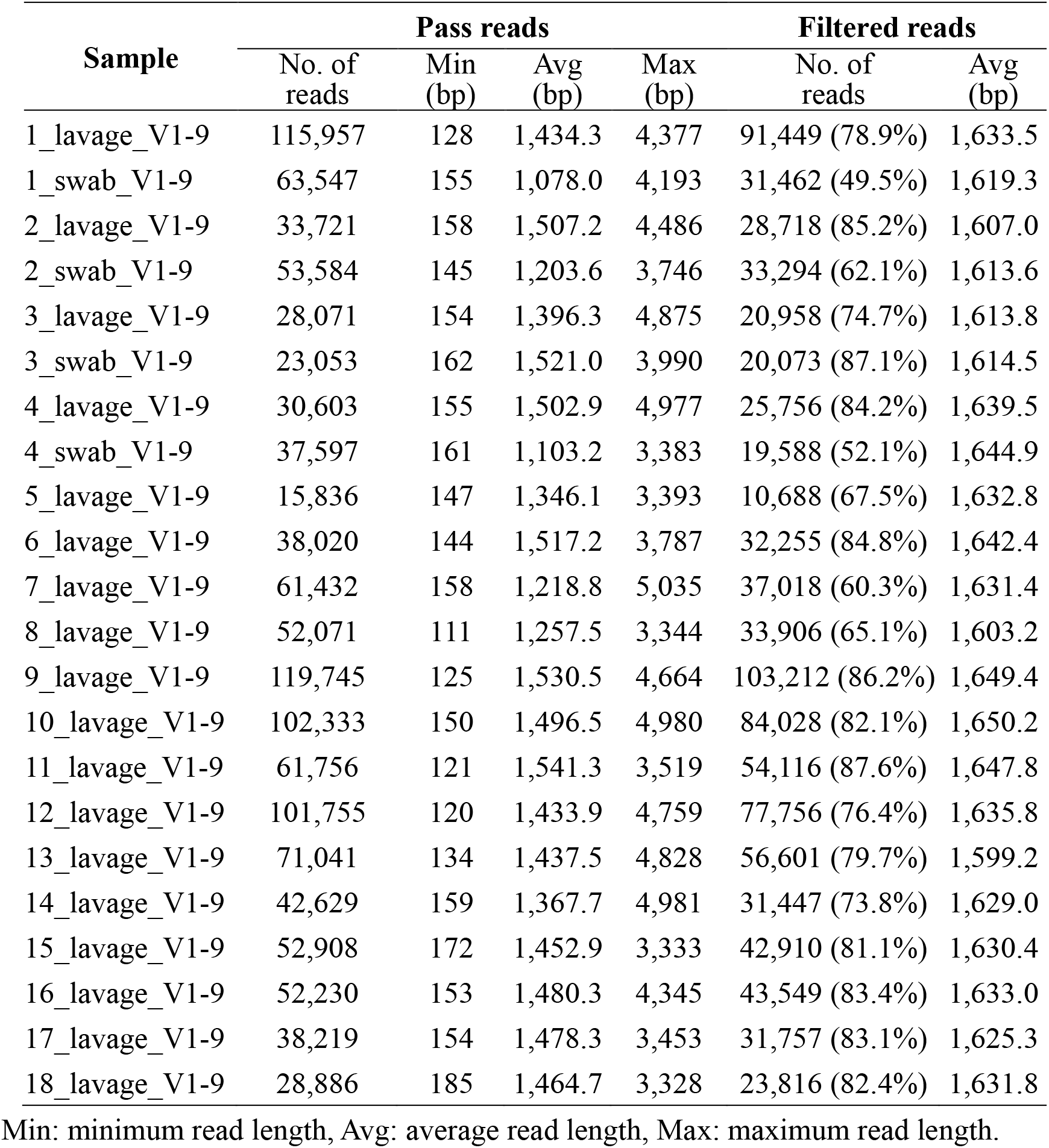
Statistics of MinION™ V1–9 sequencing data.

### Comparison of the results of swab and lavage samples

For subjects 1–4, lavage and swab samples were collected and bacterial DNA extraction and 16S rRNA analysis were performed to determine the effect of the sample collection method on the results of the microbiota analysis. After filtering, 3,000 reads were randomly extracted in eight samples (1-4_lavage and 1-4_swab), the results of which are shown in Figure 2, with almost the same identification with the two sampling methods. We performed principal coordinate analysis (PCoA) to assess the equivalence of the lavage and swab sampling methods, the results of which are shown in Figure 3. Regardless of the sample collection method, the results of the analysis of the same subjects involved almost the same coordinates.

**Figure 2.**
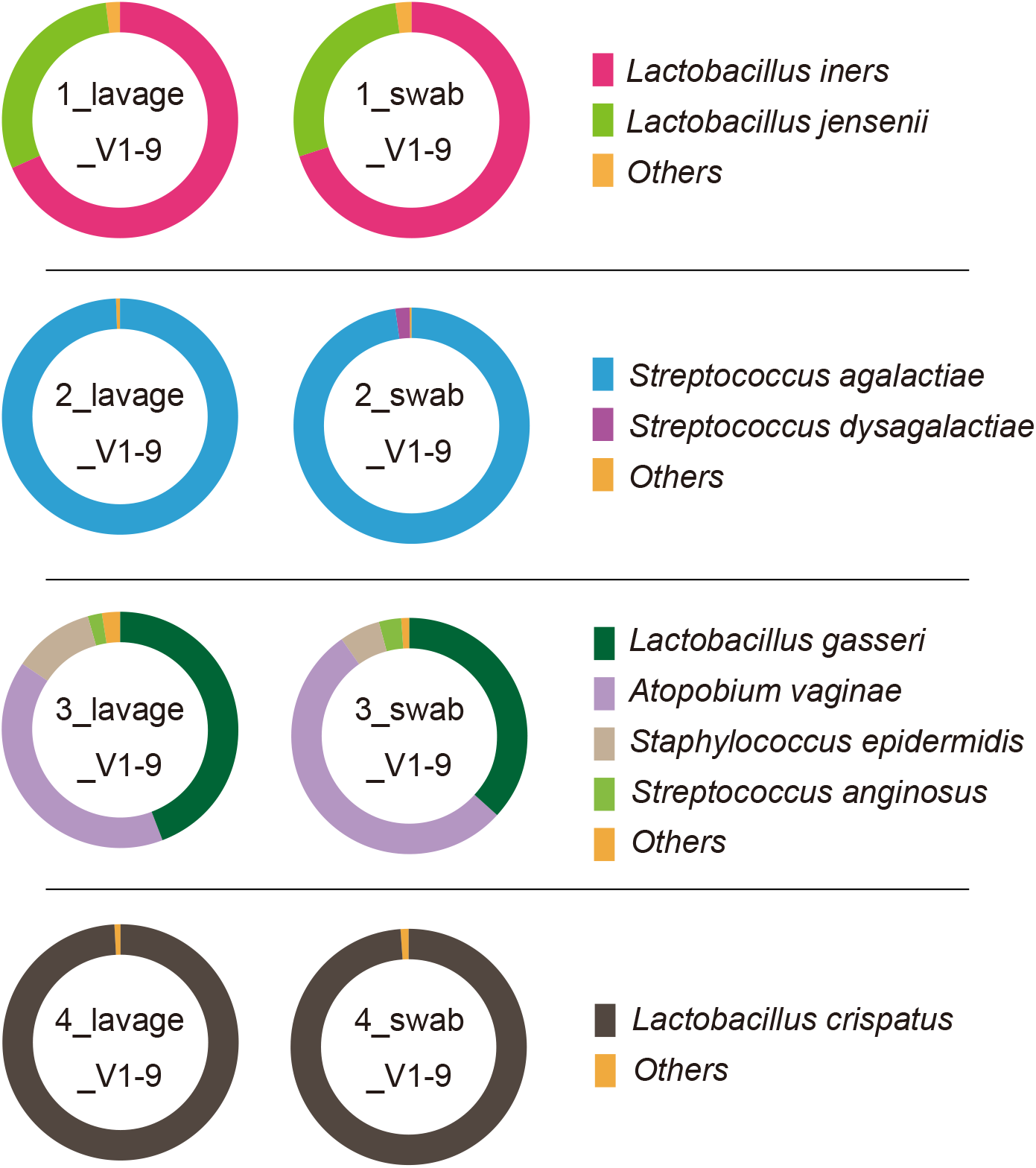
Taxonomic profiles comparing the different sampling method (lavage and swab) results of V1–9 16S rRNA MinION™ sequencing with 3,000 randomly sampled reads after filtration. In the analytical algorithm, we assigned the bacterial name that showed the highest Minimap2 score for each read; bacteria with an assignment of less than 1% were included in “Others.”

**Figure 3.**
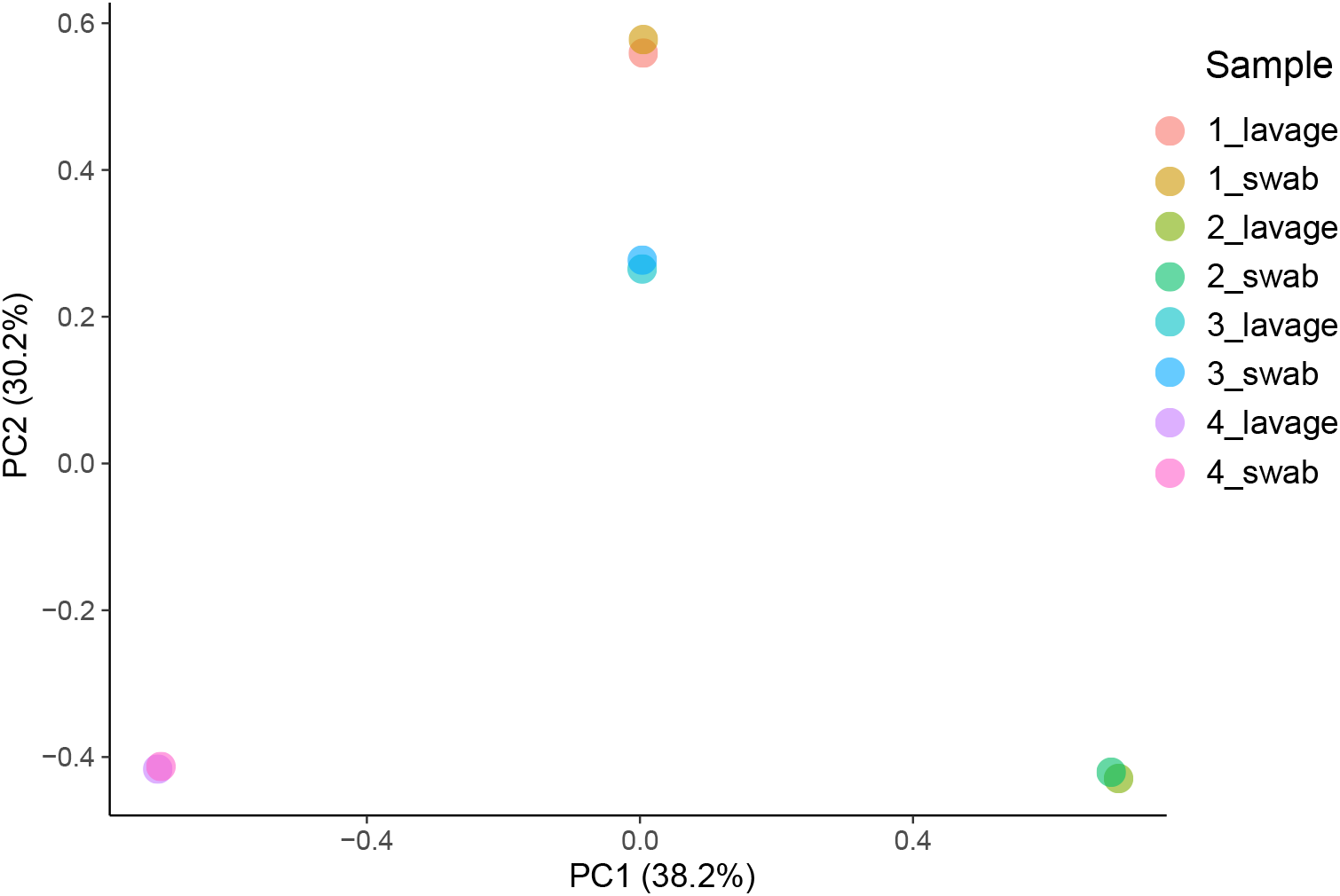
PCoA of vaginal microbiota by different sampling techniques. The coordinates of swab samples and lavage samples were nearly equivalent and did not show any effect of the vaginal specimen collection method in the 3,000 filtered reads of V1–9 MinION™ sequencing.

### Comparison of the results of sequence read number

To assess the effect of read count on the results of the MinION™ sequencing, three samples that showed characteristic bacterial microbiota [5_lavage: *Lactobacillus iners* dominant (>99%), 6_lavage: *Lactobacillus crispatus* dominant (>99%), 8_lavage: BV like] were selected and their taxonomic profiles were compared after filtering for 3,000 reads (random sampling) and 10,000 reads (random sampling) (Figure 4). In all specimens, the results of vaginal microbiota analysis in MinION™ were similar for different numbers of reads.

**Figure 4.**
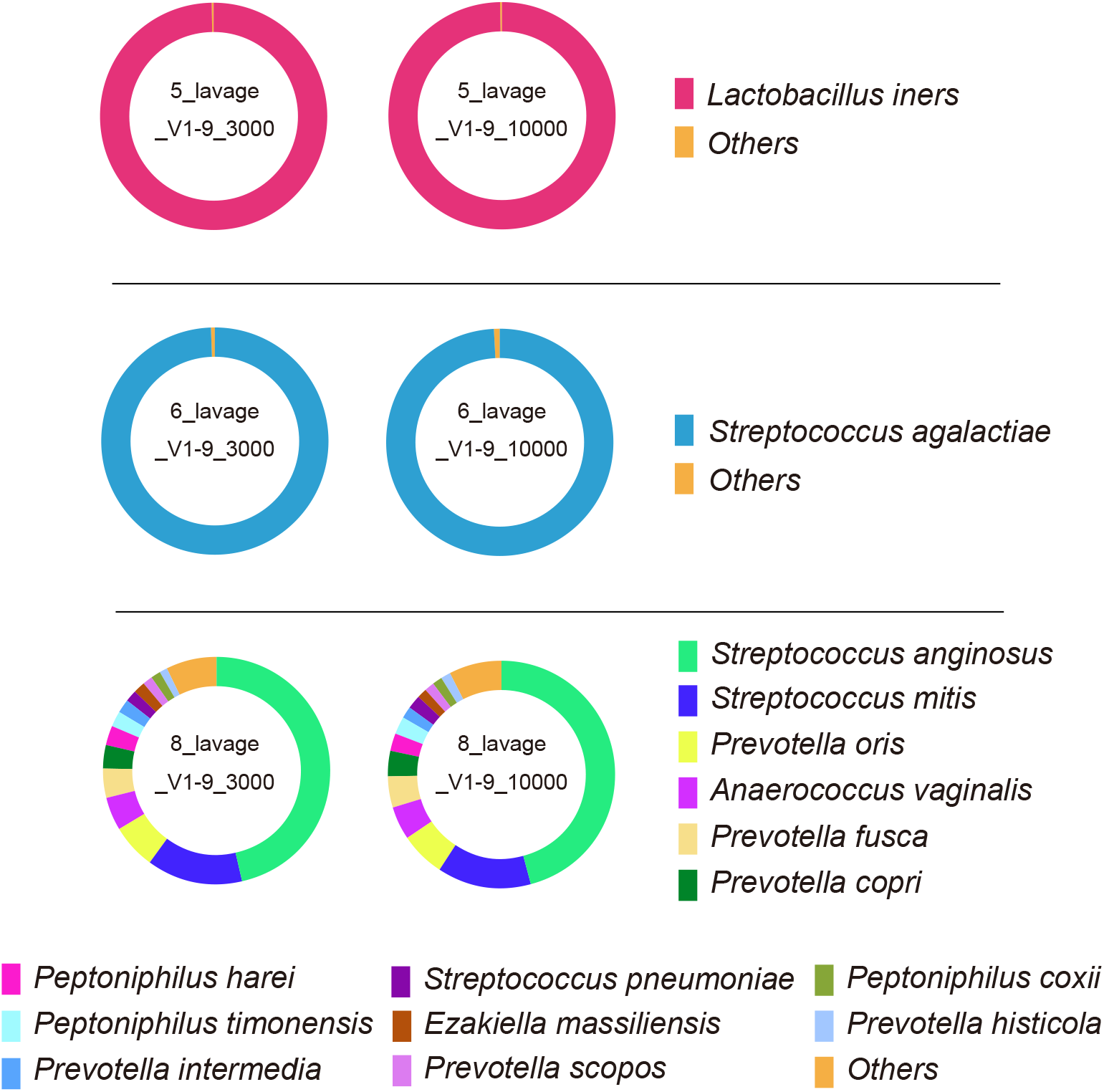
Taxonomic profiles comparing the results obtained for 3,000 and 10,000 filtered reads of V1–9 MinION™ sequencing. In the analytical algorithm, we assigned the bacterial name that showed the highest Minimap2 score for each read; bacteria with an assignment of less than 1% were included in “Others.”

### Comparison of the results of V1–9 and V3–4 16S rRNA gene sequencing

For the taxonomic classification of bacteria, we compared the resolution of long-read (V1–9) and short-read (V3–4) 16S amplicon sequencing for six samples (sample numbers 5, 6, 9, 10, 11, and 14). Of these, for sample numbers 5, 9, and 10, *Lactobacillus iners* was confirmed to be present at a rate of more than 99% (Group I), while for sample numbers 6, 11, and 14, *Lactobacillus crispatus* was confirmed to be present at more than 99% (Group C) in the V1–9 16S rRNA analysis. The V3–4 region was amplified by four-primer PCR from the bacterial DNA of six samples and sequenced with MinION™. From each sequencing result, 3,000 reads were randomly extracted and the results of the analysis of the V1–9 region were compared with those of the V3–4 region (Figure 5). In Group I, *Lactobacillus iners* was classified as being present at a rate of more than 98% in the V3–4 S16 rRNA analysis, which was comparable to the results obtained by the V1-9 sequencing (Figure 5A). In Group C, the V3-4 sequence alignment resulted in ambiguous identification of *Lactobacillus* species (Figure 5B). All samples showed *Lactobacillus crispatus, Lactobacillus acidophilus, Lactobacillus helveticus, Lactobacillus amylovorus, Lactobacillus kefiranofaciens, Lactobacillus hamsteri, Lactobacillus gallinarum, Lactobacillus kalixensis*, and *Lactobacillus psittaci* in the V3–4 S16 rRNA analysis as potentially existing species, which were found in similar proportions across all samples in Group C.

**Figure 5.**
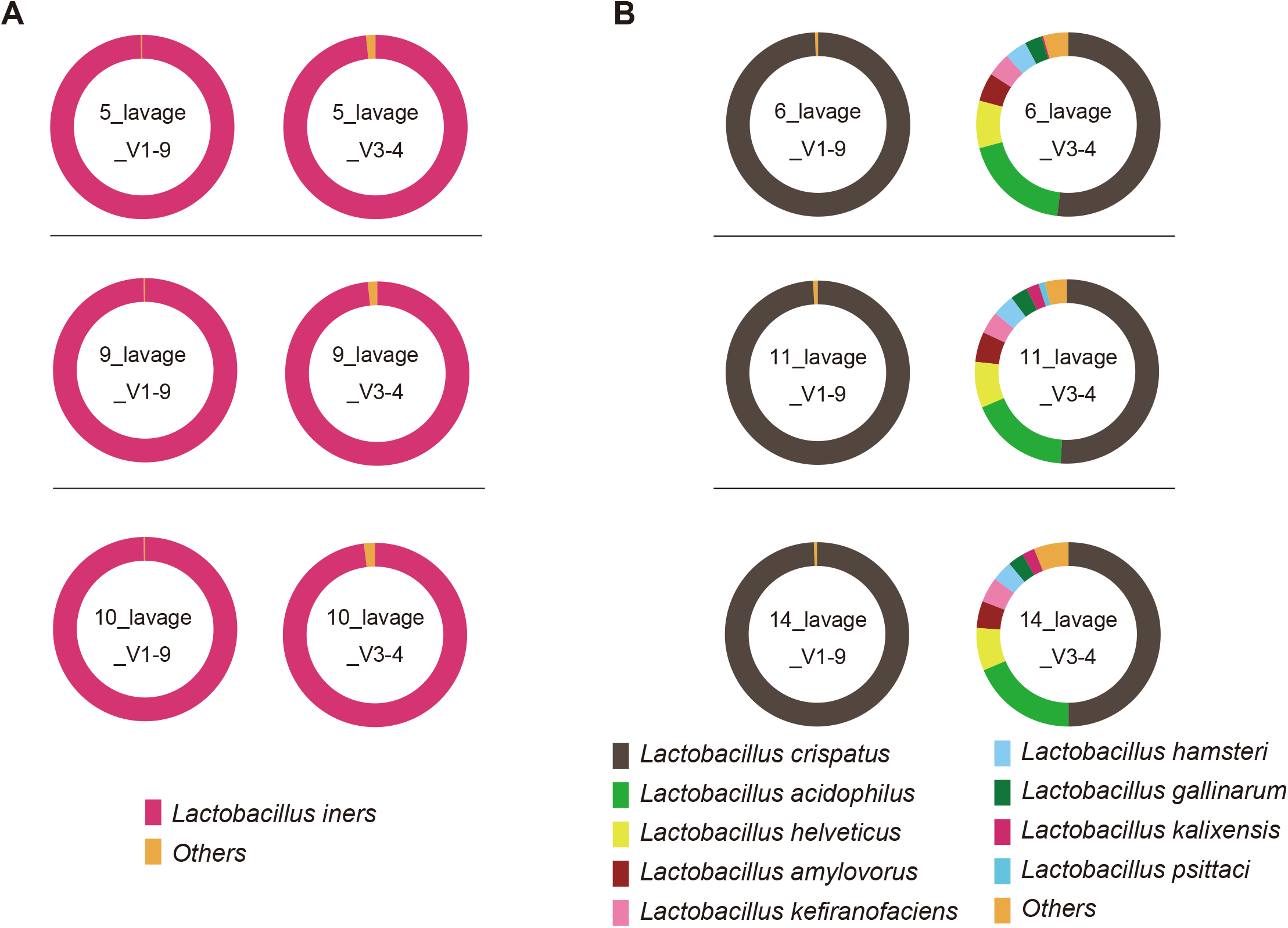
Taxonomic profiles comparing the results of V3–4 and V1–9 16S rRNA sequences using MinION™ in 3,000 filtered reads. (A) In three cases, *Lactobacillus iners* was shown to constitute more than 99% of the bacteria by MinION™ analysis with the V1–9 region (Group I). (B) In three cases, *Lactobacillus crispatus* was shown to constitute more than 99% of the bacteria by MinION™ analysis with the V1–9 region (Group C).

## Discussion

The vaginal microbiota is characterized by particularly low diversity compared with the human gut harboring a complex microbial community. [18]. Conventional methods, such as Amsel’s criteria, the Nugent score, and bacterial culture, are not suitable for an accurate bacterial identification. Next-generation sequencing has revolutionized the profiling of the bacterial microbiota. In short, the combination of metagenomic sequencing and bioinformatic technology has made it possible to more accurately assess the bacterial microbiota without the influence of uncertainties such as examiner subjectivity or the capturability of each bacterium [6]. The MinION™ platform allows a single flow cell to be used repeatedly by assigning individual barcodes. MinION™ allows data generation by immediately reading the amplicon sequences that pass through the nanopores placed on the substrate of the flow cell. The real-time data generation allows the sequencing to end when the data reach the target amount. In particular, analysis of the vaginal microbiota is a field in which MinION™’s capabilities can be maximized because of the low complexity of this microbiota under healthy conditions. However, to date, no studies have examined clinical vaginal metagenomic samples with MinION™ technology. In the present study, the number of bacterial species detected with MinION™ ranged from 1 to 22 (mean value 3.3), confirming the low diversity of the vaginal microflora (Table S2). Although no cases were diagnosed as BV by the Nugent scoring method, the 16S rRNA gene analysis using MinION™ revealed the presence of BV-causing bacteria such as *Streptococcus, Atopobium, Prevotella*, and *Gardnerella* [7] in the group categorized as intermediate based on Nugent’s criteria. For the most part, the 16S rRNA gene analysis appeared to correlate with the Nugent scoring to diagnose BV. However, there was an inconsistency between the two methods in one case (14_lavage), where it was categorized as intermediate, even though only *Lactobacillus* was identified by the MinION™ sequencing. While Nugent scoring is a subjective evaluation and depends on the acumen of the examiner, the molecular method targeting the 16S rRNA gene could serve as a practical and reliable measure for the diagnosis of BV. In vaginal microbiota analysis, it is important to be able to profile *Lactobacillus* spp. and the causative organisms of BV at the species level. The Nugent score is calculated based on the number of bacterial morphotypes and therefore has limitations for discriminating bacterial taxa at the species level. Meanwhile, the sequence-based assessment of the vaginal microbiota can allow us to obtain an accurate representation of the bacterial composition. It has been reported that species-specific characteristics of *Lactobacillus* spp. are critical for the stability and maintenance of human vaginal microbiota [19,20]. In particular, *Lactobacillus iners*-dominated community has been more associated with vaginal dysbiosis, suggesting its relevance in the pathophysiology of gynecological diseases.

Previous studies on short-read sequencing have reported that the sample collection method had no effect on the results [21]. We collected swab and lavage samples from the same cases and determined the species composition of vaginal communities. (Figure 2). When a single bacterial taxon comprised 98%–99% of the total bacteria (sample numbers 1 and 4), there was no change in the composition ratio depending on the sample collection method. When the vaginal microbiota was composed of more than two bacteria (sample numbers 2 and 3), the bacterial species comprising the microbiota were found to be the same, although there were slight differences in the proportions present. Thus, we concluded that the effect of the sample collection method was negligible. In contrast to the characterization of highly complex bacterial communities in the human gut [12], the analysis of vaginal microbiota of lower complexity would require a much smaller number of reads. A calculation using the median filtered reads predicted that 3,000 reads would be accumulated in a sequence of approximately 8 min, indicating that the time required for vaginal microbiota analysis using MinION™ could be further reduced.

The conventional parallel-type short-read sequencer cannot yield reads covering the full length of the 16S rRNA gene, and the partial 16S rRNA gene sequencing has been insufficient to identify bacterial taxa at the species level. Benchmarking with the conventional sequencing method (Illumina MiSeq™ technology) demonstrated that the full-length 16S gene sequencing by MinION™ gives a better resolution for bacterial identification [12]. To clarify the usefulness of long-read sequencing in vaginal microbiota testing, we picked up *Lactobacillus iners*-dominant (> 99% in V1–9 analysis) samples (Group I: sample numbers 5, 9, and 10) and *Lactobacillus crispatus*-dominant (>99% in V1–9 analysis) samples (Group C: sample numbers 6, 11, and 14) and compared the results of 16S rRNA analyses of the V1–9 and V3–4 regions (Figure 5). The bioinformatic pipeline used a method where each amplicon sequence was directly compared with the database, scored for similarity (Minimap2 score) according to a proprietary algorithm, and then applied to the bacterial species with the highest score. Hence, if there were multiple species with the same score, we treated the reads as potentially existing in all of the possible species. The reads assigned to “Others” in this study could be (1) bacteria with low abundance, or (2) bacteria that could not be assigned to a single species due to competition between multiple species. In Group I (Figure 5A), *Lactobacillus iners* was well recognized regardless of the choice of the regions to be sequenced, but the numbers of reads assigned to “Others” increased in the V3–4 analysis. In Group C (Figure 5B), V3–4 analysis showed that *Lactobacillus crispatus* accounted for about 50% of the reads and *Lactobacillus acidophilus, Lactobacillus helveticus*, *Lactobacillus amylovorus*, *Lactobacillus kefiranofaciens*, *Lactobacillus hamsteri*, and *Lactobacillus gallinarum* were classified in similar proportions in all three specimens, which had not been found in V1–9 analysis. Due to the lower discriminatory power of the V3-4 region, the considerable number of V3-4 reads could not be assigned to one species and evenly allocated to multiple taxonomic bins represented as potentially existing species. The numbers of reads assigned to “Others” were greater in the V3-4 data set as compared to the V1-9 data set. This suggested that assigning bacteria using the V1–9 region is more informative for determining the appropriate bacterial species than using only the V3–4 region. Similar to our previous report [12], the long-read sequencing with MinION™ showed high identification accuracy compared with the short-read sequencing under the same bioinformatic pipeline.

In terms of the usefulness of the MinION™ sequencer in gynecological practice, it is interesting that we were able to show the detailed condition of vaginal dysbiosis in a short period of time, which was not noted by the Nugent score. It has been suggested that vaginal dysbiosis may have a connection with the incidence of sexually transmitted infections [22] and genital viral infections [23,24] or cervical lesions [25]. The full-length 16S rRNA analysis using MinION™ could be a promising option for bacterial identification in a reasonable time frame for diagnostic purposes.

Having used MinION™ in our vaginal microbiota analysis workflow, we have seen four key benefits. 1) Convenience: The unit is extremely light, small, and portable, and sequencing can occur via a USB connection to an in-clinic PC. 2) Speed: The average time between specimen collection and result disclosure is 2 days. 3) Economy: Analysis of a small number of samples can be performed without increasing the unit cost of the test, by using individual barcodes. 4) Functionality: Although MinION™ has been reported to have a slightly higher error rate, the technical issues are being resolved and even the intestinal microbiota can now be identified [12]. In particular, the analysis of vaginal microbiota characterized by low diversity, as in the present study, allowed for comparable species-level identification [26]. Although this study focused on the vaginal microbiota, a similar workflow could be applied to many clinical areas in the future, and the benefits of MinION™ could make it easier for clinicians to successfully perform bacterial metagenome analysis. In addition, future large-scale microbiota studies could lead to new clinical findings.

## Conclusions

In conclusion, our validation shows that the MinION™ long-read sequencer provides a low-cost, rapid workflow for identifying vaginal microbiota with higher resolution in a clinical setting. Detecting vaginal microbiota at the species level has the potential to identify risks to women’s health, as well as facilitating large-scale clinical studies in any medical field.

## Data accessibility

Sequence data from this article have been deposited in the DDBJ DRA database (www.ddbj.nig.ac.jp/dra/index-e.html) under accession numbers DRR244979–DRR245006.

## Author contributions

Conceptualization, SK, YMa, YMo, HO, and KH; methodology, SK, YMa, SN, KK, and KH; software, SN and KK; validation, SK, YMa, and KH; investigation, SK and YMa; data curation, SK and YMa; writing—original draft preparation, SK; writing—review and editing, YMa, SN, YMo, KK, HO, and KH; visualization, SK and YMa; supervision, YMa and KH; funding acquisition, YMa, KK, and KH. All authors have read and agreed to the published version of the manuscript.

## Acknowledgments

We are grateful to Tadashi Imanishi (Tokai University School of Medicine) for his helpful comments. We would like to thank Edanz (https://en-author-services.edanzgroup.com/ac) for editing the English text of a draft of this manuscript.

## Supplementary information

Supplementary Table S1: Representative bacterial genomes stored in the GenomeSync database.

Supplementary Table S2: Details of identified species with V1-9 16S rRNA analysis using MinION™ by minimap2 best hit score (>0.5%).

Supplementary Table S3: Datils of identified taxa with V1-9 16S rRNA analysis using MinION by FASTQ 16S workflow.

## Abbreviations

BV: bacterial vaginosis
CST: community state type
PCoA: principal coordinate analysis
PCR: polymerase chain reaction
rRNA: ribosomal RNA

